# The ribosome-associated complex regulates cytosolic translation upon mitoprotein-induced stress

**DOI:** 10.1101/2025.08.21.671487

**Authors:** Jiaxin Qian, Annika Nutz, Katja Hansen, Lea Bertgen, Mirita Franz-Wachtel, Boris Macek, Johannes M. Hermann, Doron Rapaport

## Abstract

The biogenesis of mitochondria relies on the import of newly synthesized precursor proteins from the cytosol. Tom70 is a mitochondrial surface receptor which recognizes precursors and serves as an interface between mitochondrial protein import and the cytosolic proteostasis network. Mitochondrial import defects trigger a complex stress response, in which compromised protein synthesis rates are a characteristic element. The molecular interplay that connects mitochondrial (dys)function to cytosolic translation rates in yeast cells is however poorly understood. Here, we show that the deletion of the two Tom70 paralogs of yeast (*TOM70* and *TOM71*) leads to defects in mitochondrial biogenesis and slow cell growth. Surprisingly, upon heat stress, the deletion of *ZUO1*, a chaperone of the ribosome-associated complex (RAC), largely prevented the slow growth and the reduced translation rates in the *tom70Δ/tom71Δ* double deletion mutant. In contrast, the mitochondrial defects were not cured but even enhanced by *ZUO1* deletion. Our study shows that Zuo1 is a critical component in the signaling pathway that mutes protein synthesis upon mitochondrial dysfunction. We propose a novel paradigm according to which RAC serves as a stress-controlled regulatory element of the cytosolic translation machinery.

## Introduction

Mitochondrial proteins are mostly synthesized in the cytosol and then must be imported into the organelle via a set of complex processes involving multiple factors and pathways [1]. Cytosolic chaperones and their co-chaperones, particularly those from the Hsp40, Hsp70 and Hsp90 families, contribute to these pathways by maintaining mitochondrial precursor proteins in an import-competent conformation [2-5]. These (co)chaperones not only prevent aggregation and misfolding of newly synthesized proteins in the cytosol, but also ensure they are properly delivered to and inserted into the mitochondria [6]. Such interactions with (co)chaperones keep the precursor proteins in a loosely folded, unfolded, or partially folded conformation that is compatible with the import machinery [7]. The initial interactions with surface import receptors are essential for the specific recognition of the mitochondrial precursor proteins and for their subsequently efficient translocation across the mitochondrial membranes.

We and others observed that the mitochondrial surface receptors Tom70 and its paralogue Tom71 contribute to the recognition of newly synthesized mitochondrial proteins and in addition serve as a docking site for cytosolic (co)chaperones on the surface of the organelle [2-4] [8, 9]. Furthermore, Tom70 was recently reported to be involved in the localized condensation of protein aggregates on the mitochondrial surface [10]. Both Tom70 and Tom71 are anchored to the mitochondrial outer membrane via a single transmembrane segment at their N-terminal region and contain multiple tetratricopeptide repeat (TPR) domains [11, 12]. Tom70 plays a crucial role particularly in recognizing and importing precursor proteins with internal targeting sequences like multispan membrane proteins residing in both mitochondrial membranes and matrix destined proteins with internal mitochondrial targeting signal (iMTS) [13-15]. Tom71, a less abundant paralogue of Tom70, shares sequence similarity with Tom70, but its precise molecular function is still being investigated [16-18].

In line with their central role in import of mitochondrial proteins, the combined absence of both Tom70 and Tom71 causes proteostasis stress in the cytosol [2]. Such a stress manifests as accumulation of misfolded or unfolded mitochondrial precursor proteins in the cytosol, triggering cytosolic responses and potentially leading to cell damage if not resolved [19, 20]. To reduce such cytosolic proteostasis stress, cells developed a combined approach targeting protein synthesis, folding, and degradation [21-24]. Such an adaptation involves optimizing the ubiquitin-proteasome system (UPS), promoting proper protein folding, and regulating protein synthesis. The latter element is currently understudied, and it is unclear how changes in cytosolic proteostasis and mitoprotein induced stress in yeast cells can influence the quantity and quality of protein synthesis.

A potential player in translation regulation is Zuo1, also known as Zuotin, a ribosome-associated J-protein with multiple functions [25, 26]. It primarily acts as a co-chaperone for Hsp70 (Ssb1) forming together with another Hsp70 (Ssz1) the ribosome-associated complex (RAC) [27], assisting in the co-translational folding of nascent polypeptides [28]. Additionally, Zuo1 plays a role in ribosome biogenesis, DNA repair, rRNA processing, and the regulation of translation [29-31].

In the current study we investigated the interplay between the mitochondrial surface receptors Tom70/71 and the co-chaperone Zuo1. Surprisingly, we observed that, at elevated temperatures, the absence of Zuo1 improves the growth of cells deleted for Tom70/71 and accelerates translation in this double deletion strain. The absence of Zuo1 resulted also in a reduction in the levels of cytosolic stress chaperones like Hsp42 and Hsp26. Thus, we discovered a Zuo1-mediated crosstalk between accumulation of mitochondrial precursor proteins and regulation of cytosolic translation.

## Results

### The deletion of *ZUO1* has temperature-dependent effect on growth of yeast cells

To better understand the crosstalk between Zuo1 and the proteostasis of mitochondrial precursor proteins, we aimed to test the effect of the deletion of *ZUO1* on the growth of yeast cells under various conditions. To achieve that aim, we monitored growth on different carbon sources, at various temperatures, and in combination with mitoprotein-induced stress. We observed that the deletion of *ZUO1* resulted in a slower growth on fermentable carbon sources like glucose or galactose as well as on the non-fermentable carbon source glycerol (Fig. 1A-C). These results are in line with previous studies reporting a growth retardation of *zuo1Δ* cells at 30°C on glucose-containing media [25, 26]. To create cytosolic accumulation of non-imported mitochondrial proteins, we deleted the import receptor Tom70 and its less studied paralogue, Tom71. This double-deletion resulted in minorly retarded growth at 30°C on the tested carbon sources (Fig. 1A-C). Importantly, a triple deletion strain (*tom70Δ/tom71Δ/zuo1Δ)* showed a genetic negative interaction and grew much slower than the tested single or double deletion strains (Fig. 1A-C). These findings suggest that under normal temperature a functional Zuo1 is beneficial when the mitochondrial proteins Tom70/71 are absent.

**Figure 1.**
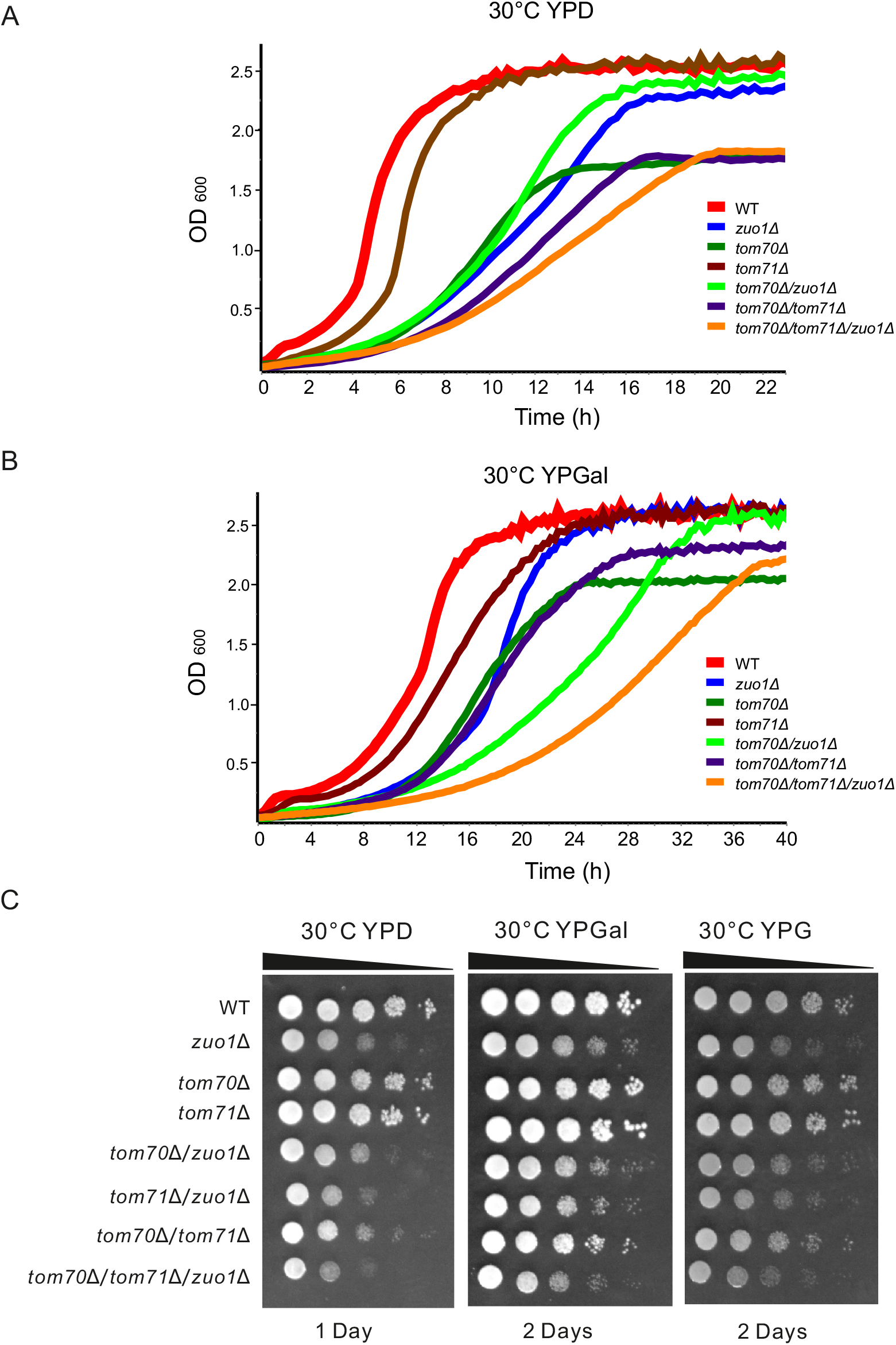
Cells deleted for *ZUO1* display retarded growth at 30°C. (**A** and **B**) The growth of the indicated strains at 30°C in rich liquid medium containing either glucose (A, YPD) or galactose (B, YPGal) was monitored using OD_600_ measurements for 24 or 40 hours, respectively. At the beginning of the measurements (time 0), the strains were diluted to an OD_600_ of 0.1. (**C**) The growth of the indicated strains was monitored at 30°C by drop dilution assay on solid rich medium containing glucose (YPD), galactose (YPGal), or glycerol (YPG). Plates were incubated for one or two days before pictures were taken.

Next, we asked whether increasing stress by growing the cells at an elevated temperature (37°C) will change the behavior of the mutated strains. As expected [2], cells lacking both import receptors hardly grew under these conditions, while the single deletion of *ZUO1* had only a mild growth defect (Fig. 2A-C). Surprisingly, the deletion of the J-protein in the strain lacking both receptors (*tom70Δ/tom71Δ)* resulted in cells that grew much better than the original double deletion strain (Fig. 2A-C). Importantly, this improvement in growth was observed on solid medium containing glucose, galactose, or glycerol as well as in liquid medium harboring either glucose or galactose. The growth enhancement upon deletion of *ZUO1* was detected also in combination with the single deletion of *TOM70* (Fig. 2A-C). In summary, at elevated temperature the absence of Zuo1 promotes the viability of cells that suffer from mitoprotein-induced stress.

**Figure 2.**
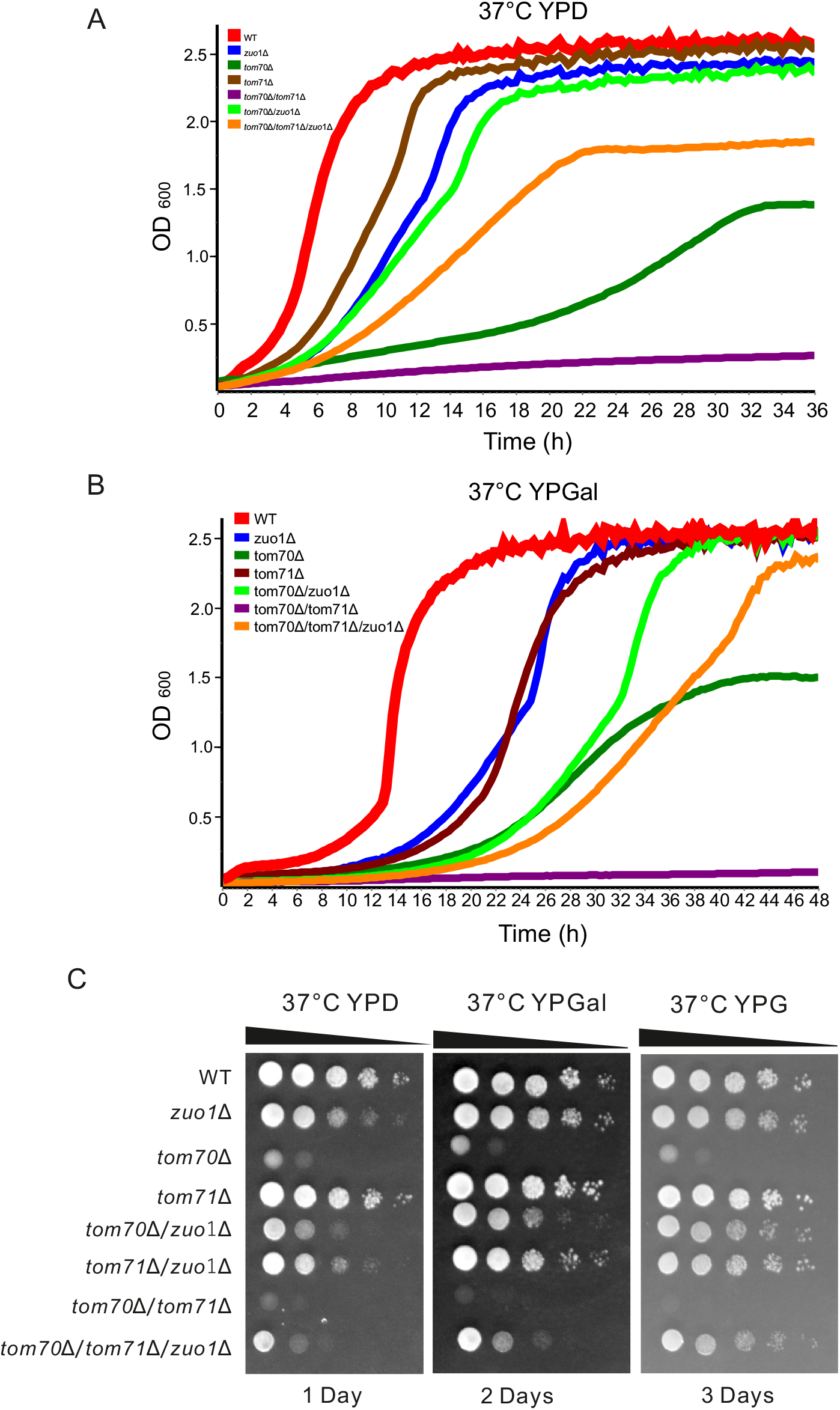
The absence of Zuo1 at elevated temperature restores the compromised growth of *tom70Δ/tom71Δ* cells. (**A** and **B**) The growth of the indicated strains at 37°C was monitored as described in Fig.1A, B. (**C**) The growth of the indicated strains was monitored at 37°C by drop dilution assay as described in Fig. 1C.

### Cells lacking Zuo1 have altered proteome

To understand the cellular events that resulted in this unexpected benefit of cells deleted for *ZUO1*, we compared by an unbiased approach the proteome of the mutated cells to those of control cells. To this goal, we isolated proteins from control (WT), double deletion (*tom70Δ/tom71Δ)*, and triple deletion cells (*tom70Δ/tom71Δ/zuo1Δ)* grown at 37°C on glucose containing medium and analyzed them by mass spectrometry (Fig. 3A). This analysis indicated a unique protein composition of the different cell types and a similar coverage of the yeast proteome among the investigated cells (Fig. 3B-C). Initially, we compared the proteome of the control cells to that of the double deletion ones. As observed before for *tom70Δ/tom71Δ* cells [2], they harbor reduced amounts of many mitochondrial proteins, among the most affected ones are Coq2 and Mdm36 (Fig. 3D-F and Suppl. Table 1). Similarly, also the triple deletion cells contained, as compared to control cells, less mitochondrial proteins where the levels of subunits of the respiratory chain complexes and mitochondrial ribosomes as well as mitochondrial inner membrane proteins were specially reduced (Fig. 3F and Suppl. Table 1).

**Figure 3.**
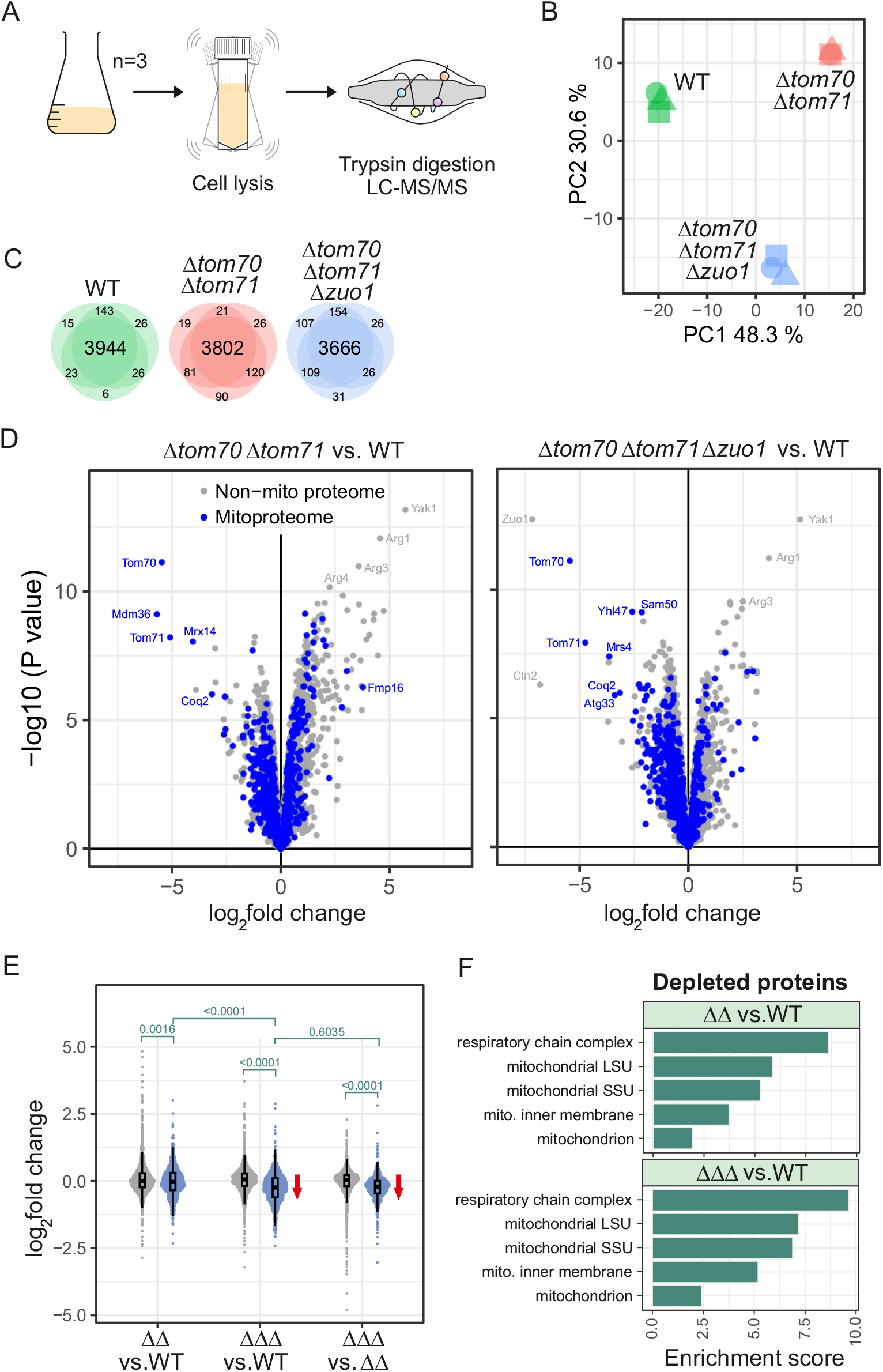
The absence of Zuo1 does not restore the depletion of mitochondrial proteins in *tom70Δ/tom71Δ* cells. (**A**) Scheme of the mass spectrometry measurement of cellular proteins. (**B**) Principal Component Analysis. (**C**) Venn diagrams showing the number of proteins that were identified in the different replicates. (**D**) Volcano plots comparing the proteomes of the indicated strains. Mitochondrial proteins are shown in blue. (**E**) Violin plots to compare the abundance of non-mitochondrial and mitochondrial (in blue) proteins in the different strains. A two-sided wilcoxon rank sum test with continuity correction was used to calculate p-values for indicated comparisons. (**F**) Enrichment of gene ontology terms was performed for the most extremely depleted proteins (log_2_ fold change > -0.8 in the limma analysis) using the GOrilla website (http://cbl-gorilla.cs.technion.ac.il); see Suppl. Table 3 for details. WT, wildtype, ΔΔ, *tom70Δ/tom71Δ*, ΔΔΔ, *tom70Δ/tom71Δ/zuo1Δ*.

A specific group of yeast genes are expressed under the control of a dedicated heat response program (Fig. 4A and Suppl. Table 2) [32]. The protein products of these genes are often involved in counteracting cytosolic proteins aggregation and misfolding. Thus, we were particularly interested in the expression of such proteins in the different examined strains. In agreement with the enhanced protein aggregation upon absence of Tom70/71, we detected elevated levels of the small chaperones Hsp12 and Hsp26 in these cells (Fig. 4B-D). Of note, comparison of the triple deletion strain with the control cells revealed that these increased expression levels were mainly diminished for Hsp26 and the levels of Hsp12 were even lower than the control cells (Fig. 4B-D). This trend was not observed for all cytosolic chaperones and the amounts of the rather generic chaperone Hsp82 were higher in the triple deletion strain as compared to control or the double deletion cells (Suppl. Table 1). Hence, our findings hint to a remodeling of the cytosolic stress response upon deletion of *ZUO1*.

**Figure 4.**
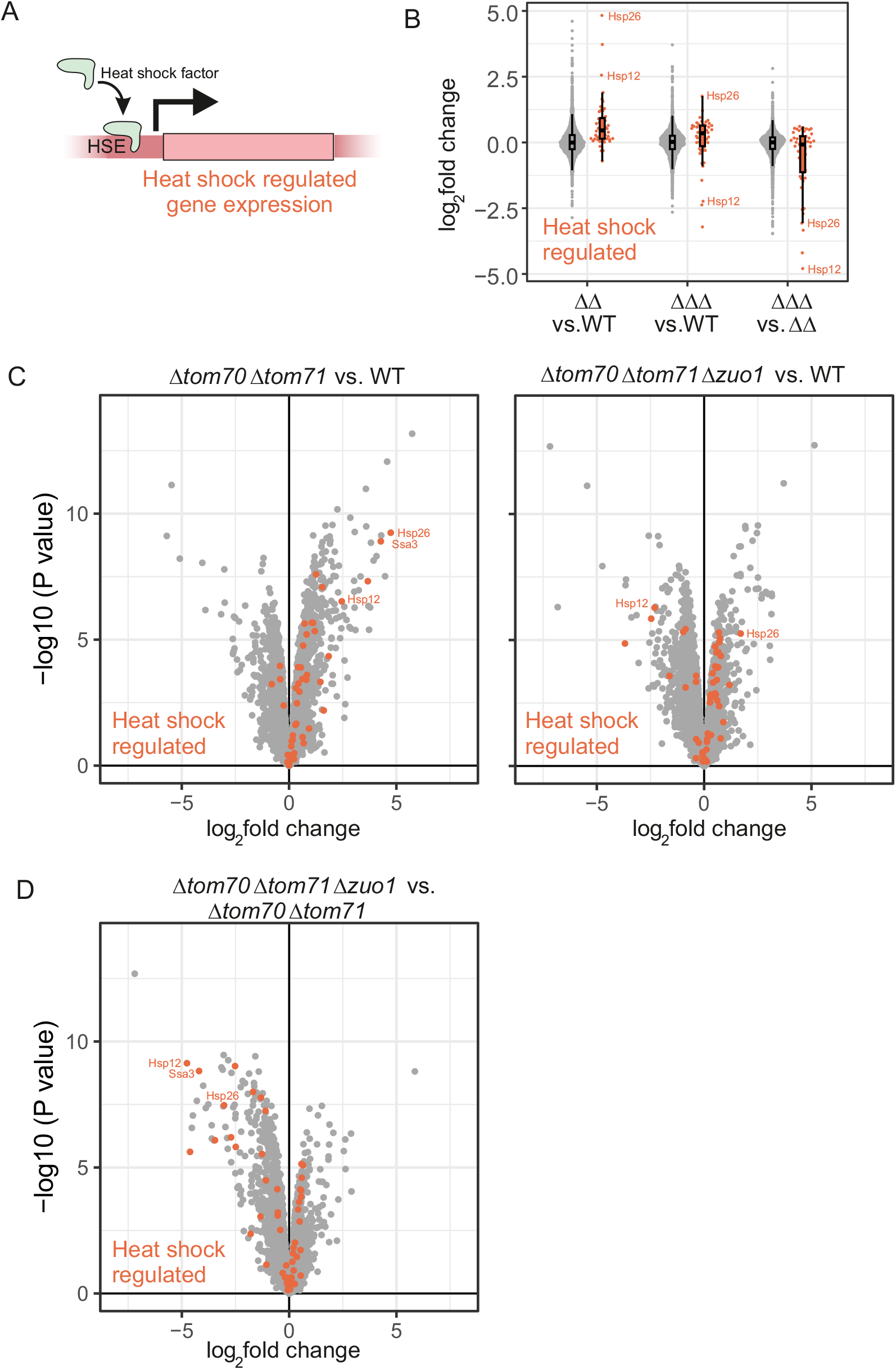
Deletion of *TOM70/71* and *ZUO1* leaves a characteristic footprint on the proteostasis network. (**A**) Scheme of the heat response program. **(B)** Violin plots comparing the abundance of proteins that are heat shock regulated (in orange, see [32] and Suppl. Table 2) to non-heat shock regulated proteins in the different strains. **(C** and **D)** Volcano plots demonstrating that deletion of *TOM70/71* leads to a marked upregulation of heat shock regulated proteins but not additional deletion of *ZUO1*. WT, wildtype, ΔΔ, *tom70Δ/tom71Δ*, ΔΔΔ, *tom70Δ/tom71Δ/zuo1Δ*.

Next, we aimed to confirm our mass spectrometry findings by an unrelated method. To this end, we analyzed the protein extracts of the investigated cells, grown on glucose at 37°C, by SDS-PAGE followed by immunostaining. Indeed, we could verify that the double deletion of *TOM70/71* caused a significant increase in the amounts of cytosolic chaperones with anti-stress activity like Hsp12, Hsp26, and Hsp104. Notably, these elevated levels were reduced back to normal levels upon the additional deletion of *ZUO1* on top of the absence of Tom70/71 (Fig. 5A-B). Interestingly, the amounts of general chaperones like Hsp70 (Ssa1) or Hsp90 (Hsp82) did not change much among the various strains. Similarly unchanged were the amounts of the unrelated cytosolic proteins Bmh1 and hexokinase (Hxk) 1 (Fig. 5A-B).

**Figure 5.**
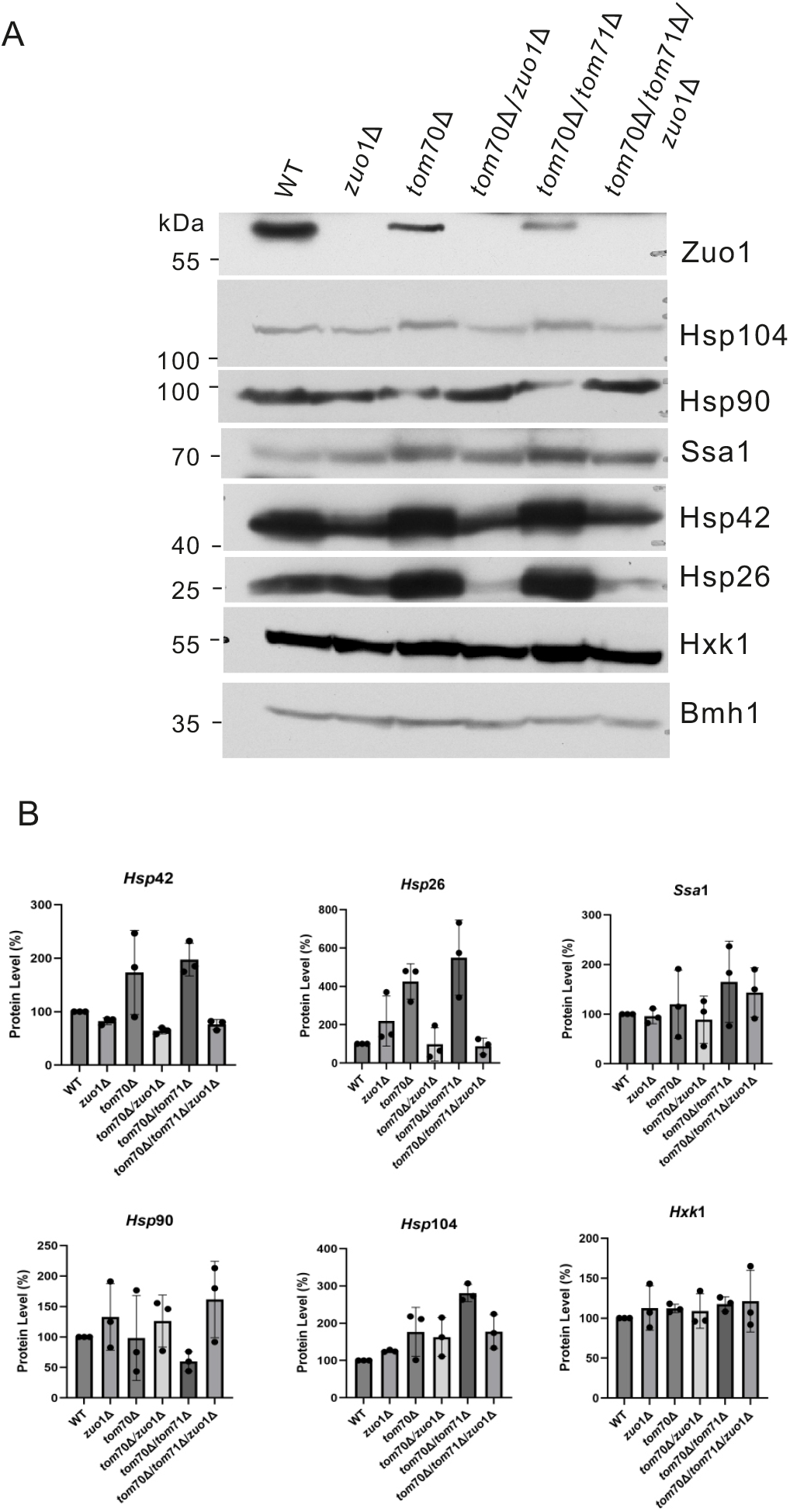
The additional deletion of *ZUO1* differentially affects the levels of cytosolic chaperones. **(A)** Proteins were extracted from the indicated strains and analyzed by SDS-PAGE and immunodecorated with the indicated antibodies. (**B**) The bands corresponding to the indicated proteins from three independent experiments as the one shown in (A) were quantified and normalized to the intensities of the Ponceau S staining. The value of the WT cells was set as 100%. The bar diagram shows the average ± SD of three independent experiments.

Since most of the mitochondrial outer membrane (MOM) proteins were reported to be substrates of Tom70/71, we also monitored their amounts in the various strains. While some proteins (like Tom20, Tom40, Msp1 and Fis1) were only mildly affected or not at all, the levels of others were dramatically altered. For example, the absence of Tom70 alone or in combination with Tom71 resulted in a major reduction in the levels of the β-barrel protein Porin (VDAC1). This reduction was partially compensated by the additional deletion of *Zuo1* (Fig. 6A-B). However, the absence of Zuo1 did not have the same effect on all proteins. A somewhat different pattern was observed for another β-barrel protein, Tob55/Sam50 where the absence of Zuo1 did not enhance the compromised levels upon the removal of Tom70 or Tom70/71 (Fig. 6A-B).

**Figure 6.**
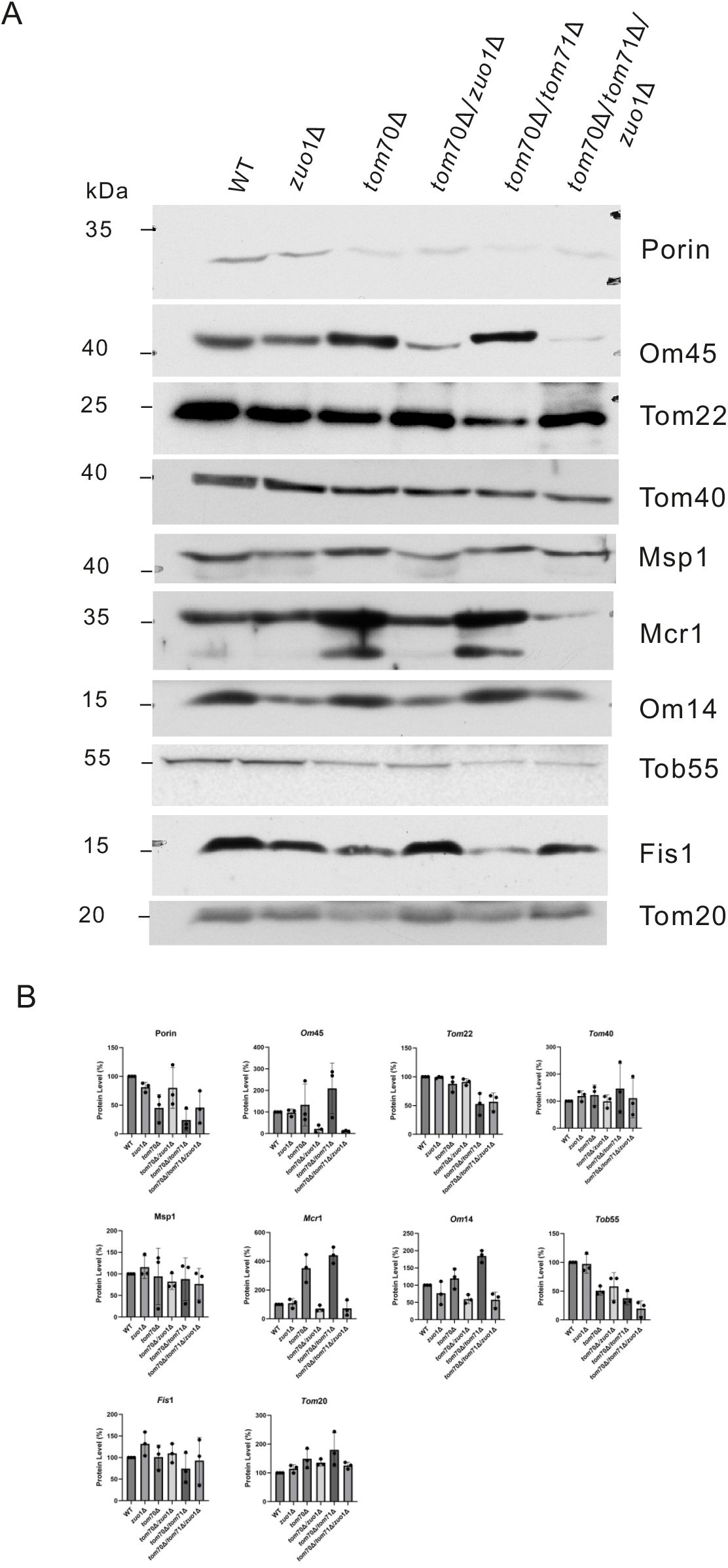
Removal of Zuo1 affects to a different extent the levels of mitochondrial outer membrane proteins. **(A)** Proteins were extracted from the indicated strains and analyzed by SDS-PAGE and immunodecorated with the indicated antibodies. (**B**) The bands corresponding to the indicated proteins from three independent experiments as the one shown in (A) were quantified and normalized to the intensities of the Ponceau S staining. The value of the WT cells was set as 100%. The bar diagram shows the average ± SD of three independent experiments.

A completely distinctive picture emerged for three proteins (Om45, Om14, and Mcr1) that were suggested to participate in mitochondrial metabolic processes. Against our expectations, the double deletion of Tom70 and Tom71 resulted in enhanced amounts of these proteins. However, the additional deletion of *ZUO1* caused a dramatic decrease in the amounts of these proteins to a level much below that in control cells (Fig. 6A-B). Taking together, the western analysis demonstrated individual effects for the absence of Zuo1 on the levels of various MOM proteins.

### The absence of Zuo1 enhances translation and changes the pattern of cytosolic aggregates

The aforementioned observations encouraged us to investigate the contribution of Zuo1 to additional cellular processes like the formation of cytosolic aggregates or translation. To monitor the number of aggregates per cell we used a previously established method namely, the staining of such aggregates with a disaggregase Hsp104 fused to GFP [33]. Using this approach, we noticed that the single absence of Zuo1 resulted in a slight increase in the number of cells with several aggregates. The double deletion of Tom70/71 resulted in even more cells with many aggregates but, the additional deletion of *ZUO1* from *tom70Δ/tom71Δ* cells reduced the number of aggregates per cell without affecting much the overall number of cells with any number of aggregates (Fig. 7A-B).

**Figure 7.**
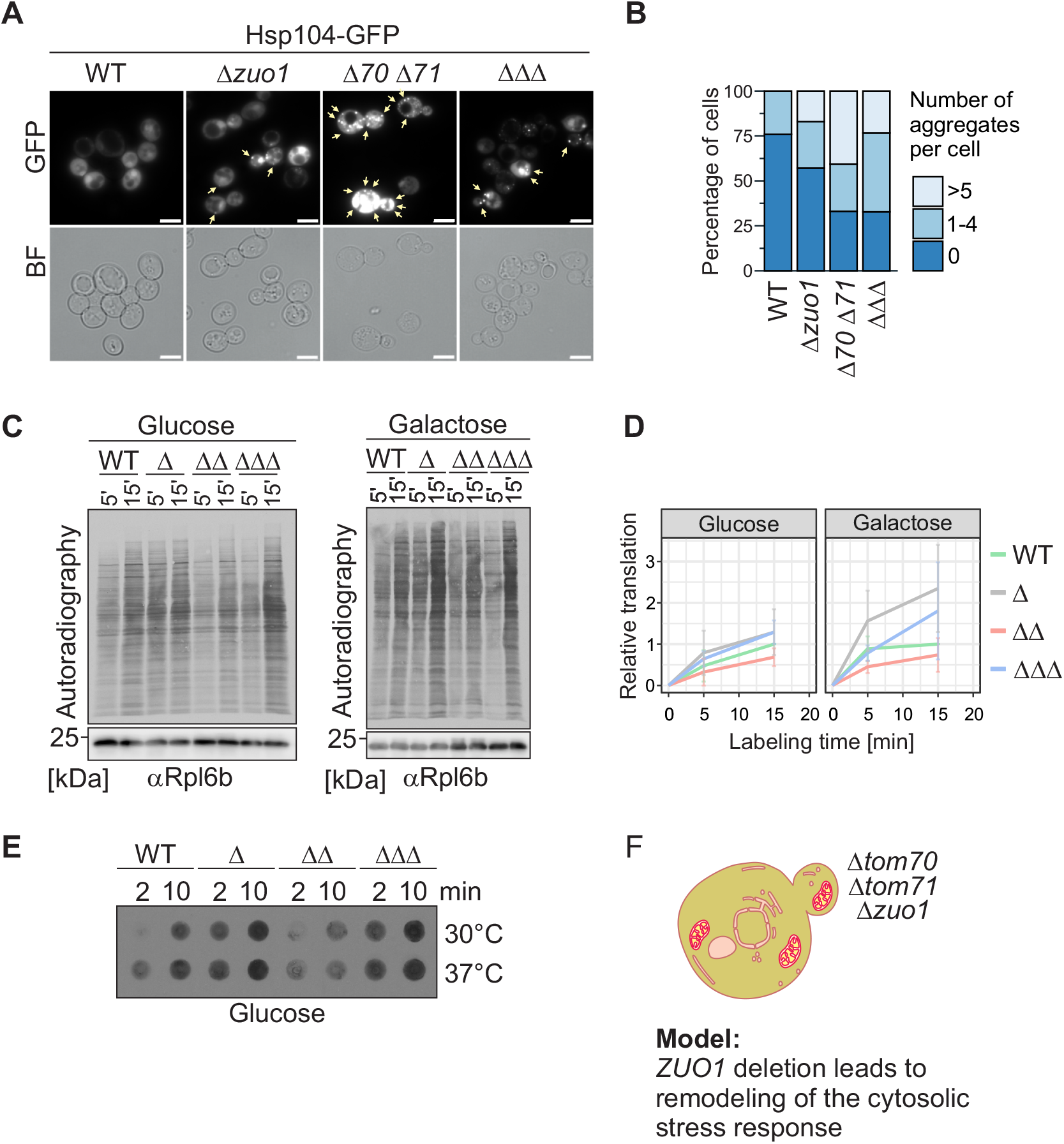
Deletion of *TOM70/71* leads to the formation of cytosolic aggregates and slows down protein synthesis. (**A**) Strains expressing Hsp104-GFP were analyzed by fluorescence microscopy. Hsp104-bound aggregates are indicated with yellow arrows. Scale bars, 5 µm. (**B**) Quantification of cells containing aggregates. Cell populations were categorized into cells with no aggregates (dark blue), with 1-4 aggregates (medium blue), or with ≥5 aggregates (light blue). The portion of cells in each category is shown as percentage of all cells counted with n = 137 for WT, n = 182 for *zuo1Δ*, n = 145 for *tom70Δ/tom71Δ*, and n = 297 for the triple mutant. (**C**) *In vivo* labeling of newly synthesized proteins at 30°C in media containing either glucose or galactose for either 5 or 15 minutes. Proteins were analyzed by SDS-PAGE and the radiolabeled translation products of the different strains were detected by autoradiography. Immunostaining against Rpl6b is shown as a loading control. (**D**) The signal of all detected proteins for a certain lane as in (C) was quantified and normalized to the loading control Rpl6b. The indicated data points at 5 and 15 min represent the relevant intensity of the radiolabeled translation products of three biological replicates. The intensity of the translation products in WT cells after 15 min was set to 1. Error bars represent ±S.D. (**E**) Cells were grown at the indicated temperature on glucose as a carbon source. Radiolabeled translation products after a labelling period of either 2 or 10 min were analyzed by dot blot and autoradiography. (**F**) Model showing that the deletion of *ZUO1* leads to the remodeling of the cytosolic stress response.

Considering this observation, we wished to determine the effect of Zuo1 on translation. To achieve this aim, we employed radioactive *in vivo* labeling followed by extracting proteins from the cells and analyzing the newly synthesized proteins by either SDS-PAGE or dot blot. Proteins were detected in both cases by autoradiography. Both assays revealed that the absence of Zuo1 increases overall translation at 30°C and the dot blot assay expanded this finding also to cells grown at 37°C (Fig. 7C-E). Collectively, these results indicate that the removal of Zuo1 from cells lacking Tom70/71 improves the growth of the latter cells probably by reducing the number of aggregates per cell in combination with enhancing global protein synthesis.

## Discussion

Defects in import of mitochondrial proteins have a well-documented impact on proteostasis in the cytosol and are associated with protein aggregation diseases [23, 24, 34]. Thus, understanding the mechanisms that contribute to proteostasis during inhibition in mitochondrial protein import can shed new light on cellular stress responses. It is well established that the absence of the mitochondrial surface proteins Tom70 and its paralogue Tom71 causes defects in mitochondrial biogenesis combined with mitoprotein stress in the cytosol [2]. This complex outcome is related to the double function of these proteins; they play a role in recognition of newly synthesized mitochondrial proteins and serve as a docking site for cytosolic (co)chaperones loaded with mitochondrial precursor proteins [3, 4, 35-37]. To better understand the cellular importance of these proteins in the context of cellular stress, we combined their absence with alteration of protein translation via deletion of the RAC subunit, Zuotin (Zuo1).

Surprisingly, at elevated temperature, these triple deleted cells (*tom70Δ/tom71Δ/zuo1Δ)* grew better than cells lacking only the two receptor proteins. This improved growth was accompanied by less aggregates per cell and enhanced translation. The latter effect is especially of interest because it is well known that the cytosolic stress response involves a coordinated effort to reduce overall protein synthesis while simultaneously promoting the synthesis of proteins essential for cell survival under stress conditions. This phenomenon, which is part of the integrated stress response (ISR), helps cells to conserve energy and resources during stress [38]. Indeed, we observed such differential alterations upon the deletion of *ZUO1*. Despite the general trend of enhanced translation, we observed that the levels of the small chaperones Hsp12 and Hsp26, which increased dramatically upon the deletion of *TOM70/71*, went down upon the removal of Zuo1.

In human cells, the link between cytosolic proteostasis stress and regulation of proteins synthesis often involves the translation initiation factor 2 subunit-α (eIF2α). It is well established that an increased level of phosphorylation of eIF2α is a hallmark of the ISR. Stress-activated kinases phosphorylate eIF2α, leading to a global reduction in protein synthesis initiation [38, 39]. Mitochondrial stress is relayed to this system in mammalian cells via the protein DELE1 (Deleted in Liver Cancer 1) [40]. When mitochondria experience stress, DELE1, which is normally imported into the mitochondrial matrix, is cleaved by the OMA1 protease and translocates to the cytoplasm, where it interacts with HRI (Heme-regulated inhibitor) kinase that subsequently phosphorylates eIF2α [40] [41]. However, despite this wealth of knowledge about the mammalian system, it was unclear how mitochondrial-induced stress is causing translation slowdown in yeast cells. Our findings shed light on this issue by suggesting that Zuo1 is part of this missing link. It seems that under stress conditions Zuo1 contributes to the reduction of translation efficiency and therefore its absence under such circumstances results in elevated protein synthesis rate (Fig. 7F).

Previous studies are not conclusive regarding the effect of Zuo1 on protein translation rate. One work reported that the deletion of Zuo1 by itself or in combination with Arsen-induced stress causes a reduction in global translation [42]. On the other hand, another report indicated that while *zuo1Δ* cells have lower polysome levels in basal conditions [43], this does not seem to affect the *in vivo* global translation rate [29]. Hence, it appears that Zuo1 might be involved in fine-tuning translation in response to different signaling inputs and in accordance with the proteostasis situation at the cytosol.

We observed that the absence of Tom70/71 causes increased number of cells with large number of aggregates and elevated levels of the small stress proteins Hsp12 and Hsp26 [44, 45], supporting their role as docking sites for substrate-associated chaperones [2]. However, the absence of Zuo1 resulted, despite the enhanced global translation, in a reduced number of cells with many aggregates and highly reduced levels of Hsp12 and Hsp26. These observations further support the notion that Zuo1 can differentially affect the synthesis of various proteins.

Our proteomic approach clearly demonstrated that the deletion of Tom70/71 resulted, as reported before, in compromised levels of many mitochondrial proteins. However, although the further deletion of *ZUO1* on this background did not restore the levels of the mitochondrial proteins, it did improve considerably the growth ability of the cells. These important findings demonstrate that the main problem in cells lacking Tom70/71 is likely not the compromised mitochondrial biogenesis but rather the cytosolic proteostasis stress. Deciphering the mechanism by which the absence of Zuo1 counteracts this latter problem is an important question that awaits future studies.

## Methods

### Growth Curves

Yeast cells were precultured in glucose medium until the exponential phase was reached. Cells equivalent to an OD_600_ of 1.0 were harvested for 1 min at 16000xg and resuspended in water. Cells were diluted to an OD_600_ of 0.1 into media containing either glucose or galactose as a carbon source. Cells were rotating at either 30°C or 37°C in a SPECTROstar^Nano^ plate reader (BMG LABTECH) with the settings shaking between reads at 500 rpm and double orbital for 20-40 hrs. Three technical replicates were used per strain and condition. The data was visualized in R.

### Monitoring Yeast Growth on solid medium

Yeast cells were initially cultured overnight in 5 mL of glucose-containing medium. The cultures were then diluted to an OD_600_ of 0.2–0.3 and incubated for further 2–4 hours until reaching an OD_600_ of 0.8–1.5. Next, cells equivalent to 2.0 units of OD_600_ were collected and resuspended in 1 mL of sterile water. A series of 1:10 serial dilutions were prepared, and 4 µL of each dilution were spotted onto agar plates with the indicated carbon source. The plates were incubated at different temperatures for further analysis.

### Whole cell lysates

Cells were cultured at either 30°C or 37°C until they reached exponential phase (OD_600_ of 0.5-1.5) and then 2-10 OD_600_ units were harvested for lysis. Cells were lysed by one of the following options: (i) Cells were washed twice with water and resuspended in 30 µl loading dye supplemented with 50 mM DTT. Cells were lysed for 5 min at 4°C with help of 4-5 glass beads (diameter of 1 mm). Afterwards, samples were boiled for 5 min at 96°C and additional 50 µl loading dye with DTT was added. (ii) Alternatively, cells pellet was resuspended in 800 µL of 0.1 M NaOH and incubated for 5 min at room temperature (RT). After centrifugation (13000 ×g, 1 min, RT), cells were resuspended in 50 µL of 2×loading dye containing 5% β-mercaptoethanol. The samples were then incubated at 95°C for 3 min, centrifuged again (13,000×g, 1 min, RT), and the supernatant was collected for further analysis. In both cases, samples were analyzed by SDS-PAGE and immunostaining. Signals were detected using the chemiluminescence mode and either the ChemiDoc MP Imaging System (BioRad) or appropriate films.

### Trichloroacetic acid (TCA) precipitation of proteins

Proteins were precipitated by adding 72% TCA to the solution to achieve a final concentration of 12% TCA. The samples were incubated overnight at -20°C to ensure complete precipitation. Proteins were sedimented by centrifugation (25,000xg, 30 min, 4°C) washed with ice-cold acetone (-20°C) and pelleted again under the same conditions. The resulting protein pellets were dried at RT and resuspended in sample buffer.

### Radioactive *in vivo* labeling of translation products

Cells were cultured in glucose- or galactose-containing medium at 30°C or 37°C until reaching the exponential growth phase and were then harvested for further analysis. For radioactive *in vivo* labeling of translation products, cells of 1 OD_600_ unit were washed with synthetic glucose- or galactose-containing medium lacking methionine. Radiolabeling was initiated by adding ^35^S-methionine (0.75 μl of an 11 μCi solution) to the suspension for incubation at the indicated temperature for 2, 5, 10, or 15 min. The reactions were quenched by adding 8 mM cold methionine in case of the dot blot experiment or cells were directly lysed after 5 or 15 min of incubation. Cells were lysed using 0.3 M NaOH, 1% β-mercaptoethanol, and 2 mM phenylmethylsulfonyl fluoride (PMSF), and proteins were precipitated with 12% TCA. The samples were analyzed using SDS-PAGE or dot blotting.

For dot blot analysis, 5 µl of a 1:30 diluted sample was loaded onto a nitrocellulose membrane using the SCR 96 Minifold I apparatus under vacuum. Radiolabeled proteins were detected by autoradiography by exposing the dried membrane to an imaging plate (Fujifilm) and visualizing it with Typhoon FLA 7000 phospho-imager (GE Healthcare) or on Super RX Medical X-Ray Films (Fuji) using the Optimax Type developer (MS Laborgeräte). Images were scanned in an 8-bit or 16-bit grayscale format, and signal intensities were quantified using Image Lab software (BioRad). Normalization was performed based on the immunostaining with antibodies against Rpl6B (antibody in a 1:2000 dilution) that served as a loading control, and the wildtype (WT) sample labeled for15 minutes was set as 100%.

### Fluorescence microscopy

Cells were transformed with a pYX142 Hsp104-GFP yeast expression plasmid [33]. Cells equivalent to one unit of OD_600_ were harvested via centrifugation, and cell pellets were resuspended in 30 µl of water. A small aliquot (3 µl) was pipetted onto a glass slide and covered with a cover slip. Manual microscopy was performed using a Leica Dmi8 Thunder Imager, with an external LED8 light source and a Leica DFC9000GTC camera. Images were acquired using an HC PL APO 100x/1.44 Oil UV objective with Immersion Oil Type A 518 F, a DFT51010 filter cube for GFP excitation at 475 nm at 540 MHz as quality mode at 16-bit and saved in the Leica Image Format. The resolution was 2048×2048 pixels and a physical size of 132.03 µm. Image analysis was performed with LAS X software, and further processing was conducted in Fiji/ImageJ. The number of Hsp104-GFP foci per cell was counted manually and categorized into three groups: cells with no foci, 1–4 foci, or ≥5 foci per cell. The percentage of cells in each category was calculated and the data were visualized using R.

### LC MS/MS

Proteins were precipitated overnight at -20°C with an ice-cold acetone-methanol mixture. After centrifugation (2000xg, 20 min, 4 °C), protein pellets were washed with 80% ice-cold acetone. Pellet was dried, and proteins resolved in denaturation buffer (6 M urea, 2 M thiourea, 10 mM Tris, pH 8.0). Protein concentration was determined with Bradford assay, and ten micrograms of protein were subjected to in-solution digestion with trypsin as described previously [46]. Desalted peptides [47] were analyzed on a Vanquish Neo nano-UHPLC coupled to an Orbitrap Exploris 480 mass spectrometer through a nano-electrospray ion source (all Thermo Scientific): 0.25 µg peptides were loaded onto a 20 cm HPLC column with 75 μm inner diameter (CoAnn Technologies, ICT36007508F-50) in-house packed with 1.9 μm ReproSil-Pur C18-AQ silica beads (Dr. Maisch HPLC GmbH, r119.aq.) under pressure control (1000 bar). Elution was performed with a 45 min segmented gradient of 5-55% of HPLC solvent B (80% acetonitrile in 0.1% formic acid) at a flow rate of 300 nl/min and a constant temperature of 40°C. The mass spectrometer was operated in a positive ion and data-independent acquisition (DIA) mode. MS1 spectra were acquired at a resolution of 120k in the range of m/z 400-1000 with automatic gain control (AGC) set to 300%, and maximum ion injection time (MaxIT) set to auto. MS2 spectra were acquired in the range of m/z 145-1450 with a normalized collision energy of 27% at a resolution of 30k, whereby AGC was set to 800%, and MaxIT was set to auto. DIA isolation windows of 10 m/z (60 scan events) were defined as the window placement optimization. The mass spectrometry proteomics data have been deposited to the ProteomeXchange Consortium via the PRIDE partner repository with the dataset identifier PXD067604.

### MS data processing

DIA-based MS data were processed in directDIA+ mode using the Spectronaut software version 19.5 (Biognosys) using default search parameters with more stringent identification settings [48]. Spectra were predicted using a *Saccharomyces cerevisiae* database obtained from Uniprot (downloaded 30th of January 2024; 6,091 entries). Results were exported using the Pivot Report showing all identified and quantified protein groups.

These were further processed using the R programming language (R version 4.5.1, https://www.R-project.org). For each condition, proteins that were identified in less than two replicates were removed (N = 3). This resulted in 4254 total identified protein groups whose quantities were log2-transformed, normalized using variance stable normalization [49], and batch-cleaned using limma [50]. Lastly, missing values were imputed by sampling N = 3 values from a normal distribution (seed = 127851) and using them whenever there were no valid values in a triplicate of a condition. The mean of this normal distribution corresponds to the 1% percentile of label-free quantification (LFQ) intensities. Its standard deviation is determined as the median of LFQ intensity sample standard deviations calculated within and then averaged over each triplicate. Proteins were tested for differential expression using limma for the indicated pairwise comparison of samples and a Benjamini-Hochberg procedure was used to account for multiple testing. All relevant test results are listed in Suppl. Table 1. Principal component analysis was carried out on the processed intensities by singular value decomposition. For gene ontology (GO) enrichment analysis of depleted proteins, proteins with smaller than −0.8 log_2_ fold change were used as target set and analyzed using the GOrilla tool (http://cbl-gorilla.cs.technion.ac.il/) [51] with all quantified proteins as background set (Suppl. Table 3).

## Conflict of interest

The authors have no conflict of interest to report.

### Author Contributions

J.Q., K.H. and L.B. planned and performed experiments and analyzed data; A.N. analyzed data; M. F.-W. performed the mass spectrometry analysis; B.M. supervised analysis; J.M.H. and D.R. planned experiments, analyzed data, and wrote the manuscript.

## Acknowledgments

We thank E. Kracker for excellent technical assistance, K.S. Dimmer for helpful discussions, and S. Rospert and J. Buchner for antibodies. This work was supported by the Deutsche Forschungsgemeinschaft (RA 1028/11-1 to D.R. and HE 2803/11-1 to J.M.H), the European Research Council (MitoCyto 101052639 to J.M.H.), and doctoral fellowship of the Chinese Scholarship Council (CSC) to J.Q.

